# High Accuracy Base Calls in Nanopore Sequencing

**DOI:** 10.1101/126680

**Authors:** Philippe Faucon, Robert Trevino, Parithi Balachandran, Kylie Standage-Beier, Xiao Wang

## Abstract

Nanopore sequencing has introduced the ability to sequence long stretches of DNA, enabling the resolution of repeating segments, or paired SNPs across long stretches of DNA. Unfortunately significant error rates >15%, introduced through systematic and random noise inhibit downstream analysis. We propose a novel method, using unsupervised learning, to correct biologically amplified reads before downstream analysis proceeds. We also demonstrate that our method has performance comparable to existing techniques without limiting the detection of repeats, or the length of the input sequence.

## I. INTRODUCTION

DNA sequencing has become a critical part of most biological research, in tasks ranging from gene network identification, to biological engineering, to organism identification and the generation of phylogenies. Most of these applications have 2 primary foci: the length of DNA sequence reads, and the perbase error in those reads. Third generation sequencing technologies presented by Oxford Nanopore Technologies(ONT) and PacBio offer read lengths 10-100x what was possible with previous sequencing technologies, but at a per-base read accuracy near 85%, down from 99.99% using second generation technologies. While the increased read length enables many new biological applications [1], [2] the low read accuracy hinders others. Extensive research has gone into developing techniques that can robustly improve Nanopore read accuracy and analyzing the trade-offs [3]–[5].

We propose an approach to correcting errors that does not make assumptions about the presence of a reference genome and is robust to mixed biological populations where reads with significant overlap must be identified as different. To accomplish this task multiple copies of the same original DNA sequence must be read, resulting in a large number reads, of which a small number are high-error copies of one another. We then use the K-means algorithm to cluster reads based on an engineered feature space. This technique does not require replicated reads to be biologically attached [6], meaning that the individual sequence lengths are not constrained. Additionally our approach groups reads with their own replicates, removing the need for a reference genome. Our contributions are summarized below:

- We demonstrate that while some base calling error is random there is a significant component that is systematic and can be modeled. in section mbox IV-A:mbox we present a simple model for predicting easily identifiable k-mers and show that our model correlates well with data found empirically.
- We propose a feature space based on the easily identifiable k-mers that accurately groups copies of unique DNA strands. In section mbox V-A:mbox, we demonstrate comparable clustering performance to MinHash [7] with a number of simulated reads similar to biological experiments.
- We demonstrate that a simple K-means approach can replace a more complex biological process to group unique DNA strands together.
- The proposed method allows for the analysis of larger DNA strands compared to INC-Seq [6].

## II. RELATED WORKS

Given the importance of sequence read accuracy, a variety of methods have been proposed for increasing read accuracy. These approaches centered around either detecting overlapping regions in genomic alignments to provide read corrections [4], [8], [9], or correcting reads before alignment [3], [6], [10], [11].

Read overlapping [9], and correction from genomic alignments [4], [8] has shown significant potential for read correction. These techniques have shown the ability to increase sequence accuracy into the high 90%’s, with relatively minimal requirements [4]. One major shortcoming is that they are limited by error rate and genomic similarity, with large genomes or high error rates resulting in excessive correction times or erroneous results [8]. Some of these shortcomings can be softened by the presence of a reasonable reference genome, as the error rate for read-genome alignments is approximately half of that seen in read-read alignments.

Read pre-correction has gained significant steam recently, and particular advantages have been seen with small genomes which contain more than 10x even within a single sequencing run. Early attempts at pre-correction focused on aligning high accuracy second generation reads to long nanopore reads [3], [11]. This provides the possibility to resolve many features but introduces systematic error within repeating regions of the reads. More recent attempts have focused on nanopore reads exclusively [6], trading a significantly reduced read length for increased read accuracy. To a large extent both of these correction efforts are unacceptable. Introducing a systematic bias into sequencing results prevents the detection of mutations in sequencing experiments, or leads to incorrect references during de novo genome constructions [3], [10]. Decreasing the relative read length is similarly detrimental in that much of the benefit of 3rd generation sequencing is consequently lost.

After correction reads typically must be mapped to a reference to be used. To aid in this mapping high-information sequences have been explored. Although because previous technologies yielded a very high level of accuracy ‘high-information’ is taken to mean rare k-mers, ones that appear less frequently than random chance would suggest. Focusing on these highly informative subsequences has allowed for significantly faster sequence matching [7], sequence pseudoalignment [12], or full alignment [13], [14]. Unlike second generation sequencing, nanopore has significantly higher error, and it is introduced as a biased error in readings. Because of this, the definition of high-information subsequences can be stretched to incorporate the probability of being able to correctly identify a k-mer.

## III. PROBLEM DESCRIPTION

DNA has a label space of Σ ∈ {*A, C, T, G*} representing the different nucleotide bases. Let *s* ∈ Σ*^n^* be a DNA strand of length *n*. Using Nanopore technology, a series of discrete measurements *δ* is generated that represents the change in electrical current as each base passes through the Nanopore from time *t*_0_ to *t*_*T*_, where *t*_*T*_ reflects the total time required for *s* to completely transit the Nanopore. This creates a corresponding vector Δ ∈ *R*^*d*^, where *d* >> *n* (*d* >> *n* (*d* ≈ *n* * 12) is the number of discrete measurements obtained from time *t*0 to *t_T_*.

The vector Δ is subsequently binned into *q*(≈ *n*) different bins, *B* = *b*_1_, *b*_2_…*b_q_*, using different time intervals that capture 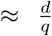 measurements per bin. The mean *μ_i_*, and standard deviation *σ_i_* are subsequently derived for each bin *b_i_*. An approximation strand, 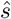, of the original strand *ŝ* is approximated from or “base called” using *μ_i_* in conjunction with a lookup table from the Nanopore manufacturer that specifies the most probable k-mer base pair for the given *μ_i_* value. Using the R9 pore from Oxford Nanopore *μ_i_* is best explained by a sequence of 6 DNA bases, thus the k-mer length *k* = 6 is used. Thus, a function that uses manufacturer provided information will yield a 6-mer *f* (*μ_i_*, σ*_i_*) = [*A*|*T*|*C*|*G*]^6^. Unfortunately, the approximation 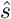 obtained from this function has > 15% [6] median error rate.

Let each approximated DNA strand be treated as a single data point 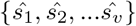, where *υ* is the total number of strands being analyzed. Let there also be ***G*** = *G*_1_, *G*_2_,… *G_p_* different groups associated with *p* unique strands. If copies are made of each strand such that *v* >> *p*, we would like to group each strand with its respective copies. The problem is formally defined as:

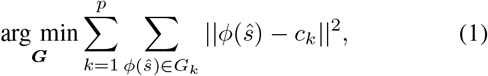
where *ϕ*(*·*) represents the transformation of a DNA strand to a descriptive feature space. *c_k_* is the centroid of group *G_k_* defined as

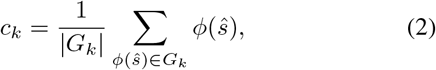
which is the mean of group *k* in the transformed feature space. Therefore, selecting an appropriate feature space transformation is vital to clustering the copies together. The objective becomes finding a feature space transformation that minimizes the distance of the centroid and its members, such that the centroid is accurately representative of each unique strand.

## IV. PROPOSED METHOD

### A. High-Accuracy K-mers

One key step in clustering DNA sequences is identifying reliable features to generate a “thumb print” for each DNA strand. This thumb print allows copies of each respective DNA strand to be grouped together with high accuracy so that subsequent sequencing corrections can be performed. In order to find the most reliable thumb print, we analyze the transition behavior of the DNA bases passing through a Nanopore. Each 6-mer DNA sequence will have six different transitions reflecting the transition of the sequence, one base at a time, through the pore. These transitions are reflected by the change in the previously defined *μ* from time *t_i_* to *t_i+1_*. Some sequences and transitions have been previously identified as being difficult to base call correctly [15]. The reference measurements provided by the nanopore manufacturer provide the mean and standard deviation associated with what one should expect to observe given a particular 6-mer DNA sequence. A state transition table is subsequently derived for each K-mer DNA base. Each transition has an associated transition value *τ* that reflects the change in value from one K-mer to the next.

We used DBSCAN [16], a density-based clustering approach, to identify the most unique *τ* patterns. The DBSCAN algorithm clusters these transitions based on density and their distance to one another. Clusters that are generated can be interpreted as sequences of k-mers which are easy to confuse with other members of the cluster, but easy to differentiate from other sequences. As a result, those clusters that tend to be smaller and more isolated reflect transition sequences that are more unique and less likely to be confused with other sequences; the more unique a sequence is, the more likely it is to be base called correctly. Because of these requirements we set the DBSCAN required density to 1 (i.e. a K-mer should be treated as a cluster even if it has no similar k-mers), and we varied the DBSCAN epsilon(neighbor euclidean distance requirement) between .5 and 5; empirically we found that an epsilon of 2 (approximately the median standard deviations provided by the nanopore the model) with k-mers of length 10 (4 transitions) gave us a sufficient number of high-quality clusters. To determine the features used we selected the 500 smallest clusters, and selected the longest common subsequence within the cluster. These k-mers were then treated as high-accuracy k-mers. A virtual DNA thumb print that is capable of clustering each unique strand with its respective copies was then generated using the high-accuracy k-mers.

The results-guided search allowed us to verify and refine our model predictions through the analysis of real-world data. To this end we gathered E. Coli K12 data previously made public by Nick Loman’s lab {http://lab.loman.net/2016/07/30/nanopore-r9-data-release/}. These reads were then aligned to the genome using Graphmap [17], while discarding unmapped reads. From the resulting alignment files the segments of exactly matching sequence were extracted, and all k-mers (for k=3-10) within them were extracted and counted. These counts were then divided by the true counts, defined by the regions on the reference genome matched by the individual reads. This ratio provides recall for all k-mers, precision is calculated by selecting all k-mers in the same range present in the reads that were not eliminated.

We find a high concordance in the k-mers identified by the model-guided and results-guided approaches, with a significant portion of the disagreement arising from subsequences with low change between dimensions as demonstrated in the first example of 2(A), where low current change between k-mers results in a higher probability of event caller error that is not accounted for in the model.

### B. Utilization of K-means

After DNA sequences have been reduced to their thumb print, the K-means algorithm was employed over the feature space to cluster DNA copies. K-means is a robust clustering algorithm that minimizes the objective function previously defined in Equation 1 while allowing us to define the number of clusters. The number of unique strands define how many clusters the K-means algorithm should find. With simulated results the exact number of unique strands is known, and future biological experiments will provide the same information (through a colony count). We find though that we are able to recover the error-corrected original sequences even in cases where the number of clusters is modestly overpredicted or underpredicted, which is of great value when the counts may not be exact due to possible sample contamination.

K-means was employed in our system using the L2 norm within the reduced subspace created by our DNA thumb print. One limitation of K-means is that it is only guaranteed to find a local minimum; to remedy this we perform 10 runs and select the run with the lowest sum of squared error (SSE). Aside from being a generic choice in K-means clustering results we find that SSE has a strong negative correlation with cluster purity on our simulated data. Finally, our implementation was employed using individual points as initial clusters; We find that this significantly removes the probability that a cluster will be empty. Additionally any cluster with less than 5 reads is treated as empty and ignored, as there is no chance of reads within being corrected to an acceptable level.

### C. Synthetic Data Generation

Current read simulators [18], [19] are incapable of properly modeling k-mer accuracy distributions identified here. nonetheless due to the low cost, high availability, and relatively high fidelity it is still valuable to validate computational techniques on simulated data. To this end we generated synthetic reads using Nanosim [19]. Each unique read was replicated according to a Poisson random value with an expected value of 50, this was empirically chosen to ensure (>99%) that at least 30 replicates were generated as we determined was required for read correction. Each replicate originates with the same sequence, but errors are calculated and applied independently. In total we generated two data sets; one with 2,000 (40 unique) synthetic reads 20,000 synthetic reads (414 unique) and performed downstream experiments on this set.

## V. RESULTS

### A. Clustering on Synthetic Data

There are not currently any standard tools for identifying replicated reads without a reference genome. Most existing tools were generated for 2nd generation sequencing technologies where reads are already high accuracy, and duplicated reads provide no additional information, and as such they are removed after alignment [20]. In this regard the closest tool we could find is the popular read overlapper MinHash Alignment Process (MHAP) [7] used for de novo genome construction. As a metric of comparison we use cluster purity as shown in figure I. To generate the following data we ran MHAP using default parameters, from the output we removed any links with less than 70% overlap, and counted connected components as clusters. In small data sets our performance is good, but less than MHAP due to MHAP performing an alignment post-processing step; in this way the full reads can be used allowing for the identification of many false positives. In larger data sets there are larger numbers of more significant overlaps that come from distinct reads.

**Fig. 1:**
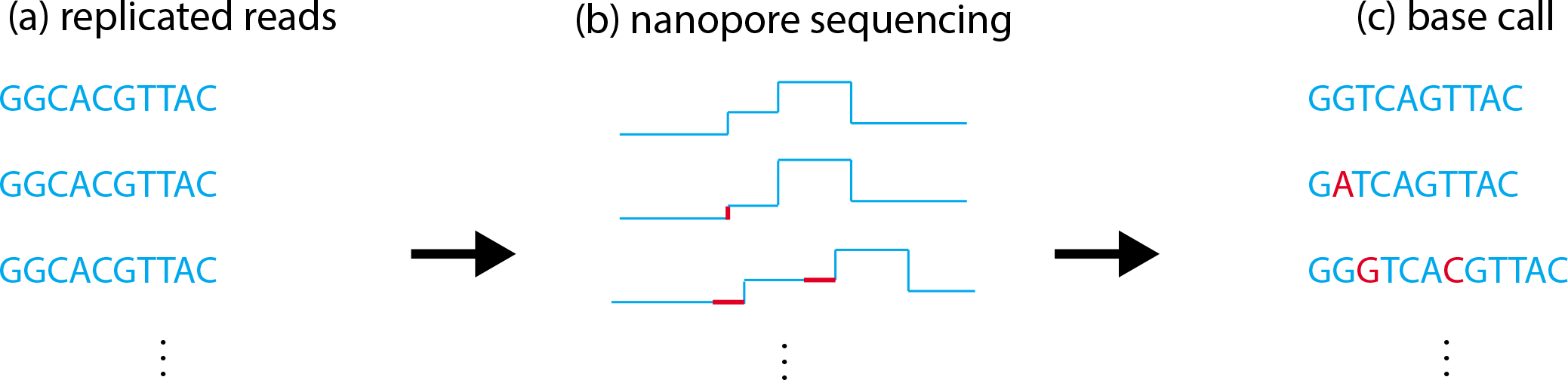
(A) Strands of DNA bases are added to the nanopore sequencer as an analog data input (B) As strands pass through the nanopores electrical current readings are taken. These readings are consequently grouped into “events”, with each event approximately representing a k-mer (sequence of bases). These events may be improperly split, events may have a mean value higher or lower than expected for a DNA k-mer, and event lengths may be different. (C) A base caller uses the sequence of events to predict the original sequence of bases.

**Fig. 2:**
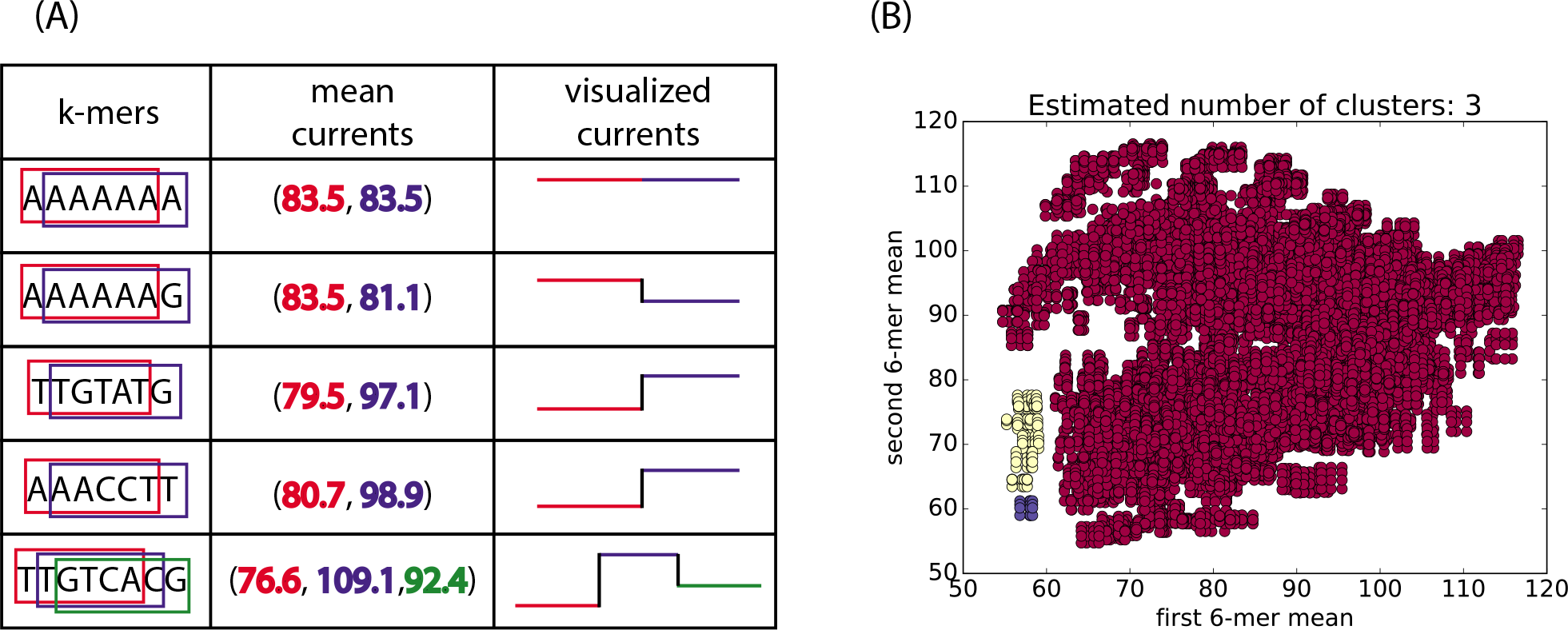
(A) DNA sequences can be interpreted as a sequence of k-mers, each k-mer can be replaced by its expected mean current to provide a mapping from DNA sequences into a high-dimensional current space. (B) Clusters of 2-dimensional points (sequence length 7), clusters are found using DBSCAN with neighborhood size 1 and epsilon 2

**Table 1:** Comparison of MHAP and K-means clusters over the engineered feature space, the number of K-means clusters was set to the number of components identified by MHAP

| Organism | Simulator | number of reads | MHAP purity | K-Means purity |
| --- | --- | --- | --- | --- |
| E. Coli | NanoSim | 2,000 | 98.2% | 96.6% |
| E. Coli | NanoSim | 20,000 | 77.2% | 75.9% |

Despite the utilization of cluster purity as a metric, the ultimate goal is to generate accurate reads to be used for downstream processing. To this end we utilized PBDAGCON [10] to create consensus sequences from aligning reads within each cluster. We find that in the smaller E. Coli data set we are able to accurately recover each of the 40 unique reads with a mean read accuracy of 97%, up from an individual mean read accuracy of 81%. In the larger data set we are able to recover 408 of 414 individual sequences, again with a mean accuracy of 97%. On inspection most of the failing reads are caused by a division of read copies between multiple clusters. Interestingly, while some sequences can be base called with only 15 copies, depending on the error distribution the consensus caller can fail with as many as 30 copies, indicating that biological sequencing experiments should aim to have more than 30 copies for read correction.

## VI. CONCLUSION

We have proposed a novel unsupervised learning technique capable of significantly increasing accuracy of nanopore base calls. We demonstrate that our technique provides significant advantages over current techniques by providing similar levels of accuracy while removing read length limitations imposed by previous techniques. We tested our method on data simulated from a state of the art simulator, and hypothesize that on actual results the performance should be even better.

The availability of high accuracy reads allows for the exploration of new applications, including; sequencing of larger organisms, organism disambiguation when sequencing a population, exact sequence detection in diploid and polyploid organisms, and the ability to scaffold genomes across exceptionally long repeat regions. Providing a path to these high accuracy-reads without compromising read length is therefore crucial to continued progress.

